# Identification of general acid catalyst for the ATPase activity of Lynch syndrome-related MutL homologs

**DOI:** 10.1101/2021.07.09.451751

**Authors:** Kenji Fukui, Yuki Fujii, Takato Yano

## Abstract

Mutations of mismatch repair MutL homologs are causative of a hereditary cancer, Lynch syndrome. Investigation of MutL facilitates genetic diagnoses essential for cancer risk managements and therapies. We characterized MutL homologs from human and a hyperthermophile, *Aquifex aeolicus*, (aqMutL) to reveal the catalytic mechanism for the ATPase activity. Although existence of a general acid catalyst had not been conceived in the mechanism, analysis of the pH dependence of the aqMutL ATPase activity revealed that the reaction is accelerated by general acid-base catalysis. Analyses of mutant aqMutLs showed that Lys79 is the general acid, and the corresponding residues were confirmed to be critical for activities of human homologs, on the basis of which a catalytic mechanism for MutL ATPase is proposed. These and other results described here would contribute to evaluating the pathogenicity of Lynch syndrome-associated missense mutations.

## Introduction

DNA mismatch repair (MMR) is a highly conserved DNA repair system, which corrects mismatched bases that are generated by DNA replication errors, recombination between non-identical sequences, and other biological processes (1-5). MutS homologs initiate MMR by recognizing mismatched bases and subsequently recruit MutL homologs (5,6). In the majority of organisms excluding *Escherichia coli* and other γ-proteobacteria, MutL homologs have an endonuclease activity to incise the newly-synthesized, i.e. error-containing, strand of the mismatched DNA duplex (7-9). The mismatched base on the incised strand is removed by exonucleases and the resulting gapped region is filled by DNA replication polymerases.

Since MMR significantly contributes to the genome integrity, mutations and epigenetic silencing of the genes encoding human MutS and MutL homologs sometimes cause Lynch syndrome that is one of the most major hereditary cancers (10-12). Therefore, genetic diagnosis of MMR genes is attracting a great attention in the view of cancer risk management.

Genetic diagnosis of MMR genes is also critical for selection of particular cancer therapies. The MMR pathway participates in induction of apoptosis that is caused by alkylating and cross-linking agents (13). Therefore, MMR-deficient cancer cells are resistant against anticancer drugs such as *N*-methyl-*N*’-nitro-*N*-nitrosoguanidine and cisplatin (14-17). In addition, it was recently found that MMR-deficient solid tumors are extremely sensitive to immune checkpoint blockade with antibodies to programmed death receptor-1 (18,19). This is because the MMR-deficient cancer cells contain high numbers of somatic mutations and develop a large number of immunogenic neoantigens that are targeted by the immunotherapeutic response (19,20). Thus, genetic diagnosis of MMR genes can be utilized to predict responses of tumors to these anticancer therapeutics and help planning the therapy strategies.

In the past decade, the development of next-generation sequencing technologies enabled identification of numerous mutations of the MMR genes from Lynch syndrome-suspected patients. However, a large amount of those mutations are so-called variants of uncertain significance (VUSs), and it is difficult to determine whether the mutations are pathogenic or benign (21). Presence of VUSs prevents the rapid and accurate genetic diagnosis of MMR genes. Abundance of VUSs in MMR genes is due to the lack of understanding on the function of the MMR proteins at the amino acid residue level. Identification of functionally-important amino acid residues in the MMR proteins is an urgent task to deal with MMR-associated cancers.

MutL homologs belong to the GHKL ATPase/kinase superfamily that includes DNA <u>g</u>yrase B subunit (GyrB), <u>H</u>sp90, bacterial histidine <u>k</u>inase, and mitochondrial serine <u>k</u>inase, in addition to Mut<u>L</u> (22-24). The GHKL superfamily proteins generally consist of the N-terminal ATPase and C-terminal dimerization domains. The N-terminal domain (NTD) contains a motif comprising four highly conserved amino acid sequences, the Bergerat ATP-binding fold (Fig. 1) (24). ATP binding and hydrolysis at the Bergerat ATP-binding fold induce large conformational changes, which are essential for molecular function of the GHKL superfamily proteins.

**Figure 1.**
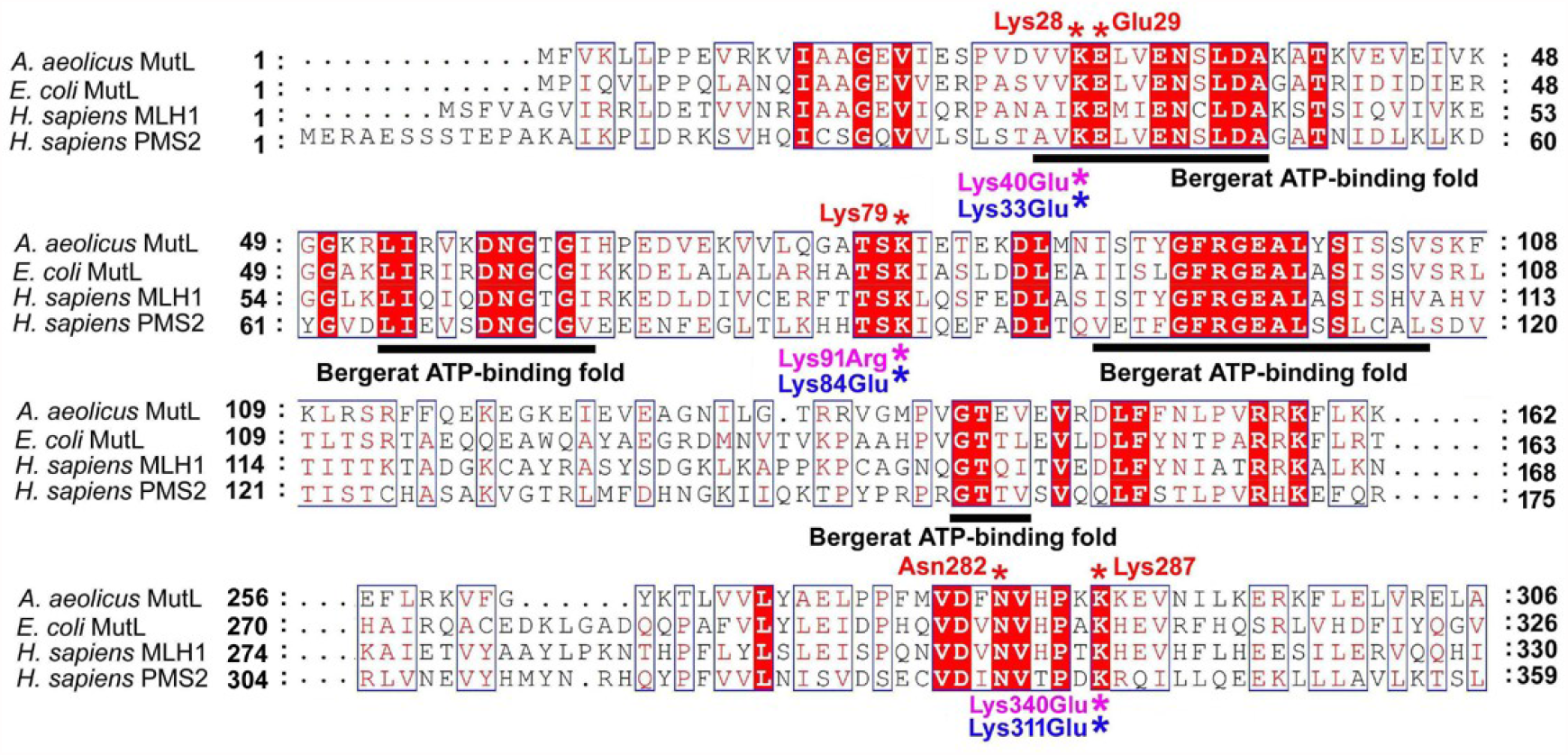
Amino acid sequence alignment of MutL homologs. The amino acid sequences of the ATPase domains of aqMutL, *E. coli* MutL, human MLH1, and human PMS2 were aligned by CLUSTALW (25) and visualized using the ESPript3 program (26). Regions corresponding to residues 163–255 of aqMutL are omitted for simplification. Pink and Blue asterisks indicate Lynch syndrome-associated mutations clinically found in the human *PMS2* and *MLH1* genes, respectively. Red asterisks indicate the residues of aqMutL that were mutagenized in this study. Black bars under the sequences indicate the Bergerat ATP-binding fold.

Bacterial MutL endonucleases are homodimeric, whereas eukaryotic counterparts are heterodimeric. Human MutLα is comprised of MutL homolog 1 (MLH1) and postmeiotic segregation increased 2 (PMS2) and is receiving increased clinical attention because a significant number of Lynch syndrome-associated mutations are found in the genes encoding MLH1 and PMS2. Although MLH1 and PMS2 show significant amino acid sequence similarity to each other, only PMS2 contains the metal-binding motif that is responsible for the endonuclease activity and is conserved in MutL homologs from a wide range of organisms including most of bacteria. The endonuclease activity of the MutL C-terminal domain is thought to be regulated by the ATP binding- and hydrolysis-induced conformational changes (27-29), and the ATPase activity of MutL proteins has been shown to be critical for *in vivo* MMR activity (30,31).

The catalytic mechanism for the ATPase activity of MutL proteins has been learned from structural and functional analyses on *E. coli* MutL (23), eukaryotic MLH1 (32), and PMS2 (33). In the crystal structure of the *E. coli* MutL NTD complexed with an ATP analog (23), Glu29 coordinates a water molecule that is located at a position suitable to attack the γ-phosphoryl group of the substrate. Replacement of Glu29 by an alanine resulted in a complete loss of the activity (23). Glu34 of human MLH1 and Glu41 of human PMS2, corresponding to Glu29 of *E. coli* MutL, have also been characterized to be essential for the ATPase activity (32). On the basis of these results, Glu29 of *E. coli* MutL is thought to be the general base to activate the nucleophilic water. In addition, it was demonstrated that the K307A mutation in *E. coli* MutL resulted in a 20-fold decrease in the ATPase activity (23). Since Lys307 is located near the γ-phosphoryl group of the substrate, it has been thought that Lys307 also participates in the catalysis. However, the catalytic mechanism for the ATPase activity of MutL remains to be elucidated.

Previously, crystal structures of the NTDs of *E. coli* MutL (23,34), *Aquifex aeolicus* MutL (aqMutL) (35,36), human PMS2 (33), human MLH1 (37), and yeast PMS1 (a counterpart of human PMS2) (38) were solved. Although the tertiary structures of these proteins are highly similar to each other, several differences are found in details of their structures. The largest difference is found in the linkage between ATP binding and dimerization of the domains. The binding of ATP analogs to the *E. coli* MutL and human MLH1 NTDs induces the transformation of five unstructured loops into ordered structures that serve as the dimerization interface. In contrast, in the human PMS2 and aqMutL NTDs, binding of ATP analogs does not induce the transformation of the unstructured loop regions; hence, the ATP analog-bound forms of these proteins are monomeric. Thus, aqMutL exhibits similarity to human PMS2 and is thought to be suitable as a model molecule to study the molecular mechanism for the ATPase activity of human PMS2 (36).

*A. aeolicus* is a hyperthermophilic eubacterium that grows at temperatures over 90°C (39). aqMutL and other proteins from this bacterium are extremely stable against temperature and pH changes and, therefore, suitable for physicochemical characterizations. In this study, we examined the pH dependence of the aqMutL ATPase activity, which led to identification of the catalytically important lysine residues. One of the lysine residues was found to be the general acid in the enzymatic reaction. It was confirmed that the corresponding lysine residues are also critical for the *in vitro* ATPase activities of human MutL homologs and the *in vivo* MMR activity in the eubacterium *Thermus thermophilus*. Based on these results, we propose a model for the catalytic mechanism of the MutL ATPase activity.

## Results and Discussion

### pH dependence of the aqMutL ATPase activity

Generally, plots of reaction rates of an enzyme against pH reflect ionization states of amino acid residues that are directly involved in the catalysis. Although proteins from human and other mesophilic organisms are generally not tolerant against extreme pHs, it has been known that proteins from thermophilic microorganisms are highly stable under a wide range of pH conditions (40,41). Therefore, we used aqMutL as a model molecule to study the catalytic mechanism of the MutL ATPase activity. Prior to examining the pH dependence of the aqMutL ATPase activity, we tested the pH stability of the full-length aqMutL. aqMutL was incubated at given pHs for 3 h at 60°C, and the circular dichroism (CD) spectra were measured (Fig. S1). The negative peak values at 222 nm were constant over a pH range from 3.5 to 10.7 (Fig. 2A), indicating that aqMutL was stable under these pH conditions.

**Supplementary Figure S1.**
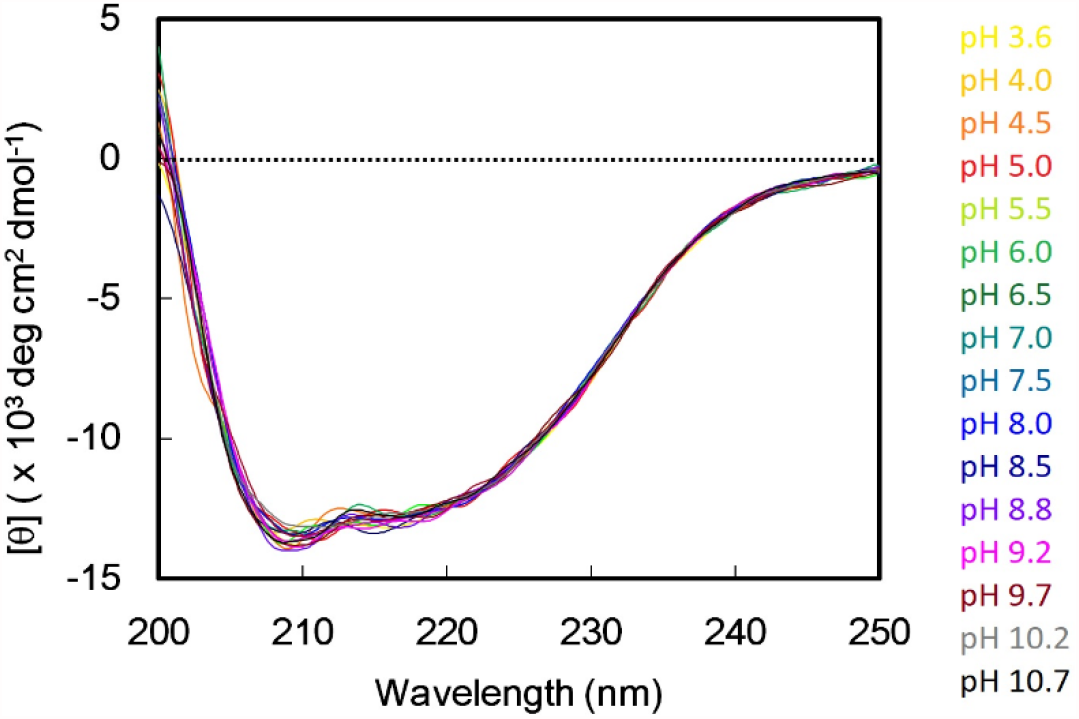
Far-UV CD spectra of aqMutL under various pH conditions. aqMutL was incubated in a pH range 3.5–10.7 for 3 h at 60°C, and the CD spectra were measured. Each spectrum was generated by accumulating wavelength-scan data 10 times.

**Figure 2.**
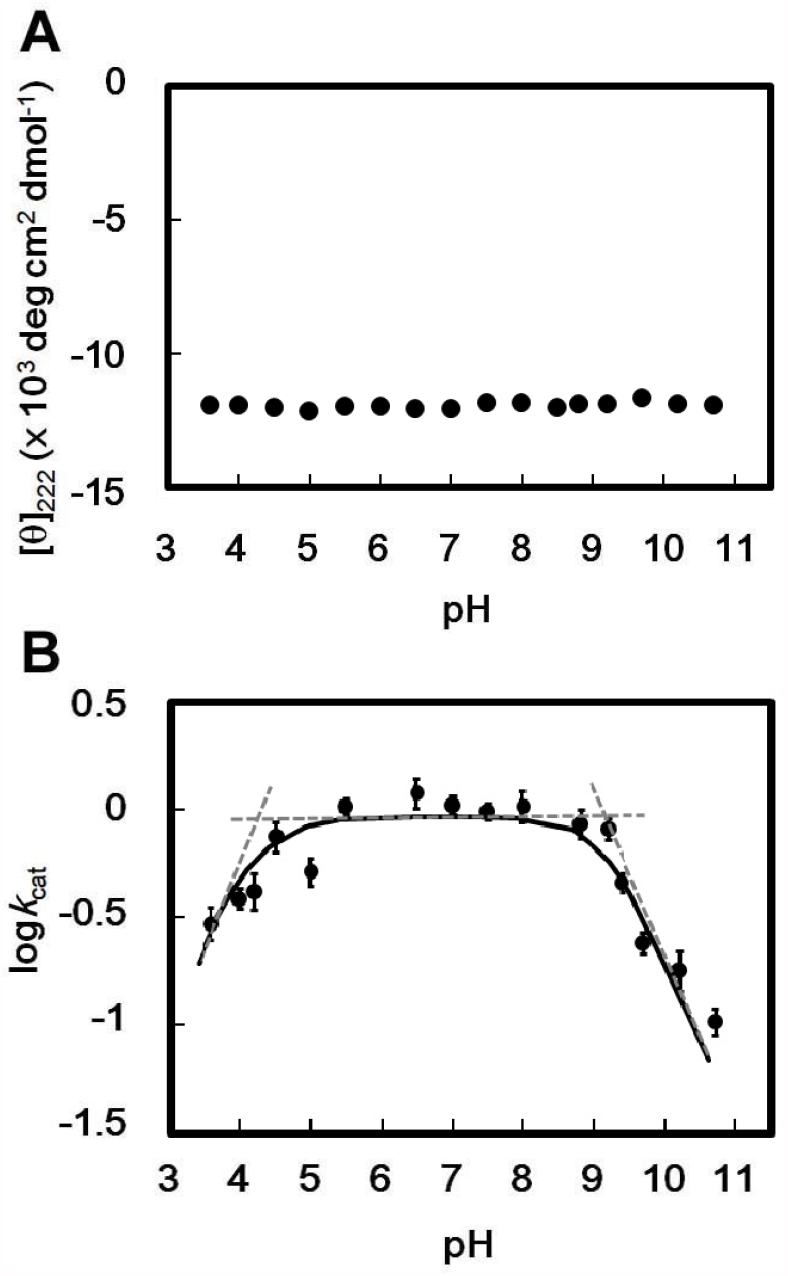
Effects of pH on the stability and ATPase activity of aqMutL. (A) aqMutL was incubated in a pH range 3.5–10.7 for 16 h, and the CD spectra were measured. The residue molar ellipticity values at 222 nm were plotted against pH. (B) The ATPase activity of aqMutL was measured in the same pH range, and the log*k*_cat_ values were plotted against pH. *k*_cat_ values are turnover numbers per minute. Experiments were repeated three times, and bars indicate the standard deviations. A theoretical line derived from the scheme described in the method section is shown as a solid line. Broken lines are drawn to visually indicate the two p*K*_a_ values, 4.1 and 9.4, obtained by the analysis.

The ATPase activity of aqMutL was measured by detecting the release of the product phosphate using a colorimetric assay. The assay was performed at various ATP concentrations in a pH range from 3.5 to 10.7, and each *k*_cat_ value was determined by fitting the standard Michaelis-Menten equation to the data. The *k*_cat_ versus pH profile was bell-shaped with maximal *k*_cat_ values at pH 5–8 (Fig. 2B). The profile is best explained by that the enzyme, or, to be more precise, the enzyme-substrate complex, has two catalytically essential dissociable groups; one with a p*K*_a_ of 4.1 has to be deprotonated and the other with a p*K*_a_ of 9.4 protonated for the catalysis to proceed.

It is known that free ATP exhibits two p*K*_a_ values, around 4.3 and 6.5, which reflect dissociation of protons from *N*_1_ of the adenine ring and the γ-phosphoryl group, respectively. Previous studies, however, indicated that Glu29 of *E. coli* MutL, corresponding to Glu29 of aqMutL, serves as the general base in the catalysis (23). Therefore, we postulated that the bel-shaped profile of the pH dependence reflects the protonation states of the enzyme aqMutL. The existence of a general acid with an alkaline p*K*_a_ has not been postulated in the MutL reaction, and, thus, we aimed to identify an amino acid residue functioning as the general acid catalyst.

### Lys28 is important for the protein stability and involved in the ATPase activity, but not the general acid catalyst

In the crystal structure of the ATP analog-bound form of the aqMutL NTD (36), Glu29 of the Bergerat ATP-binding fold (Fig. 1) forms a hydrogen bond with a water molecule that is located at an optimal position to attack the γ-phosphoryl group of ATP (Fig. 3A). This is consistent with the notion that Glu29 functions as the general base to activate the nucleophilic water. Interestingly, the ε-amino group of Lys28 interacts with the carboxy group of Glu29. Lys28 is included in the Bergerat ATP-binding fold and conserved in all MutL homologs (Fig 1). To the best of our knowledge, no previous report has studied the role of this lysine residue. In addition, substitution of a glutamate for Lys40 in human PMS2 or Lys33 in human MLH1, which corresponds to Lys28 of aqMutL, has been found in Lynch syndrome-suspected patients and classified as a VUS (Fig. 1).

**Figure 3.**
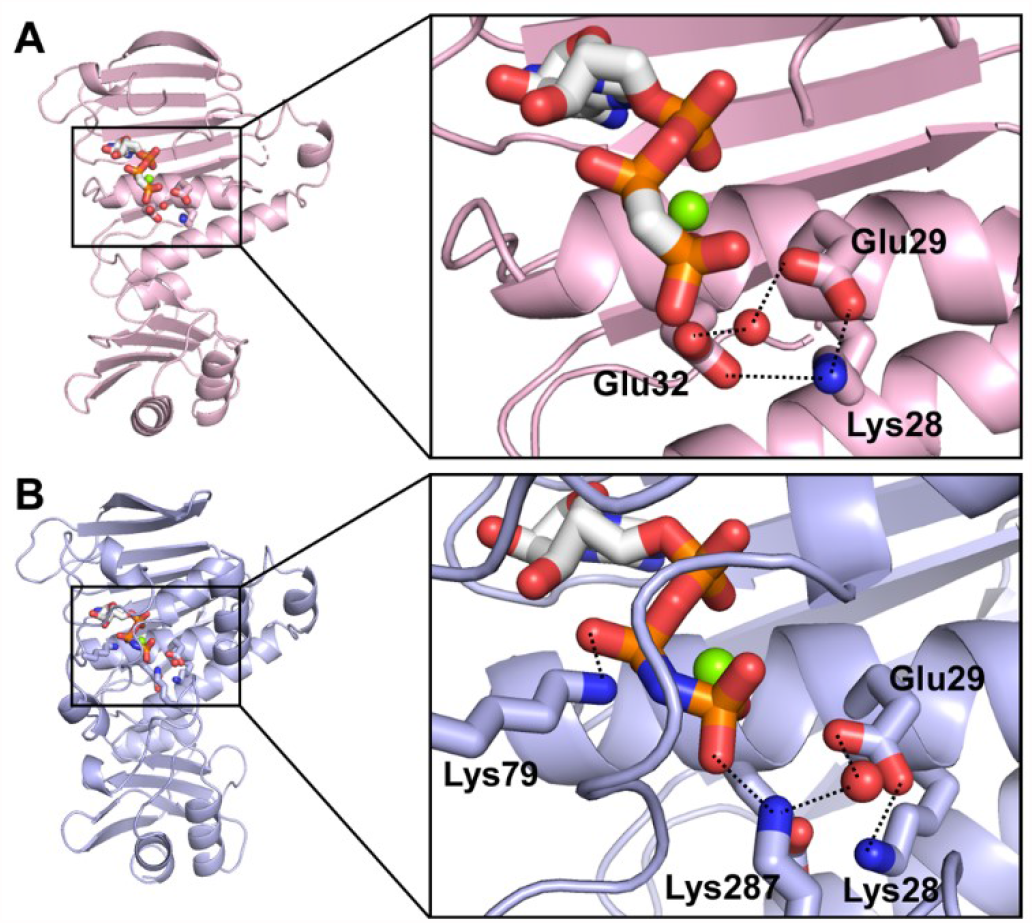
ATP-binding site of the aqMutL NTD. (A) The crystal structure of the aqMutL NTD complexed with the ATP analog AMPPCP (PDB ID: 6LZJ). (B) A model structure of an ATP analog-bound state of the aqMutL NTD was generated by the SWISS-MODEL homology-modeling pipeline (42) using the crystal structure of the AMPPNP-bound *E. coli* MutL NTD (23) as a template structure. Magnesium ion and the nucleophilic water are shown as green and red spheres, respectively. The ATP analogs and the side chains of the interacting residues are shown in stick models. In the model structure, Lys79 and Lys287 interact with the β- and γ-phosphoryl groups of the ATP analog, respectively.

We created the K28E aqMutL and examined its physicochemical properties and ATPase activity. The CD spectrum of the K28E aqMutL was exactly the same as that of the wildtype aqMutL (Fig. 4A), indicating that the mutation does not influence the overall structure of the protein. However, the CD intensity of the K28E aqMutL started to decrease at lower temperatures and lower concentrations of a denaturing agent than the wildtype aqMutL (Fig. 4B and C). These results suggest that the K28E mutation destabilizes the structure of aqMutL.

**Figure 4.**
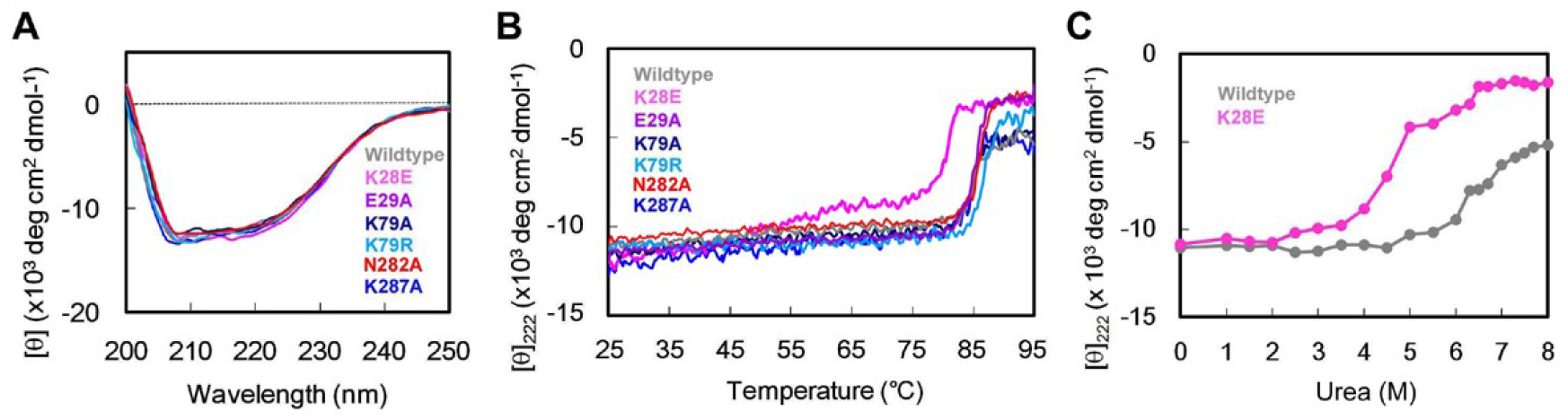
CD measurements of the wildtype and mutant aqMutLs. (A) Far-UV CD spectra. Each spectrum was generated by accumulating wavelength-scan data 10 times. (B) Temperature-dependent denaturation was monitored by measuring the residue molar ellipticity at 222 nm. (C) Urea-dependent denaturation of the wildtype and the K28E aqMutLs. The residue molar ellipticities at 222 nm were plotted against urea concentrations (0–8 M).

The ATPase activity of the K28E aqMutL was measured, and the kinetic parameters were determined (Fig. 5A and Table 1). Although the *k*_cat_ value was decreased by 3.4 fold, the K28E aqMutL retained significant activity. Moreover, the pH dependence of the *k*_cat_ values still exhibited the p*K*_a_ value, 9.2, of the general acid catalyst (Fig. 5B). These results indicate that Lys28 is involved, but not essential, in the catalysis and that another residue serves as the general acid for the catalysis.

**Table 1.** Kinetic parameters for the ATPase activity of aqMutLs, ProS2-tagged human PMS2 NTDs, histidine-tagged human MLH1 NTDs, and *A. aeolicus* GyrB NTDs.

| Proteins | $k_{\text{cat}}$ ($\text{min}^{-1}$)<sup>1</sup> | $K_{\text{m}}$ (mM)<sup>1</sup> |
| --- | --- | --- |
| aqMutL WT | $1.5 \pm 0.15$ | $0.36 \pm 0.19$ |
| K28E | $0.44 \pm 0.022$ | $0.90 \pm 0.073$ |
| E29A | N.D.<sup>2</sup> | N.D.<sup>2</sup> |
| K79A | N.D.<sup>2</sup> | N.D.<sup>2</sup> |
| K79R | $0.17 \pm 0.014$ | $0.61 \pm 0.13$ |
| N282A | $0.029 \pm 0.0010$ | $0.56 \pm 0.054$ |
| K287A | $0.46 \pm 0.023$ | $0.37 \pm 0.036$ |
| PMS2 WT | $0.37 \pm 0.018$ | $0.19 \pm 0.039$ |
| K91A | N.D.<sup>2</sup> | N.D.<sup>2</sup> |
| K91R | $0.044 \pm 0.0017$ | $0.30 \pm 0.042$ |
| K340A | $0.063 \pm 0.0081$ | $0.16 \pm 0.093$ |
| K340E | $0.029 \pm 0.0018$ | $0.080 \pm 0.012$ |
| MLH1 WT | $0.16 \pm 0.0065$ | $0.79 \pm 0.075$ |
| K84A | N.D.<sup>2</sup> | N.D.<sup>2</sup> |
| K84E | N.D.<sup>2</sup> | N.D.<sup>2</sup> |
| K311E | $0.020 \pm 0.0030$ | $0.75 \pm 0.13$ |
| aqGyrB WT | $2.2 \pm 0.13$ | $0.25 \pm 0.041$ |
| K109A | N.D.<sup>2</sup> | N.D.<sup>2</sup> |

<sup>1</sup>The standard Michaelis-Menten equation was fitted to the data to determine the $k_{\text{cat}}$ and $K_{\text{m}}$ values.
<sup>2</sup>N.D. means that the activity was not detected.

**Figure 5.**
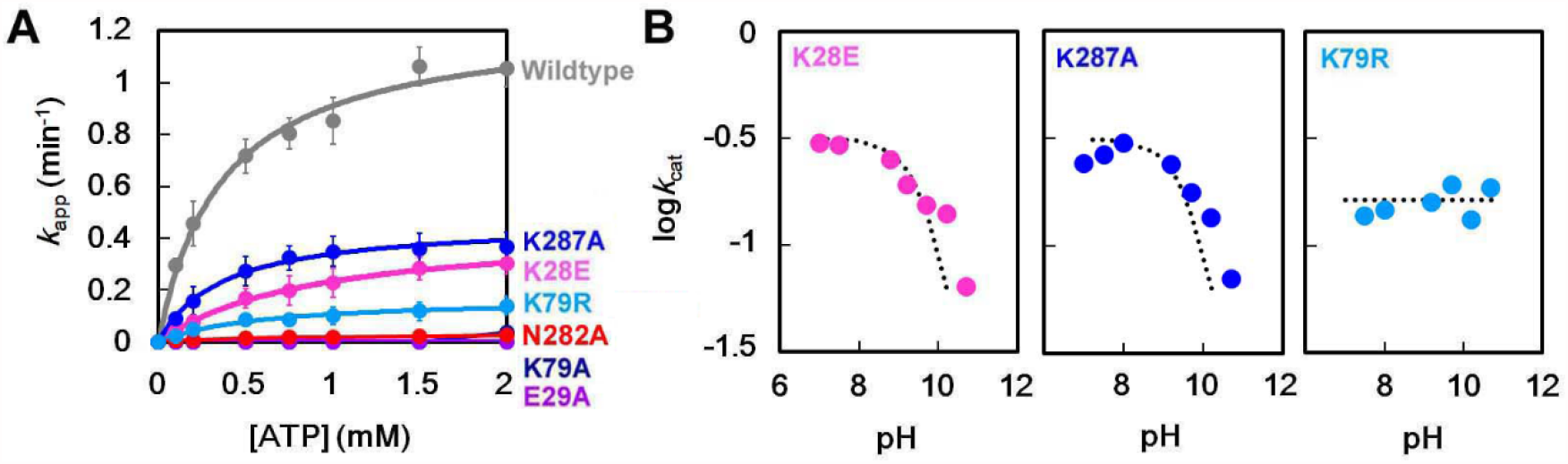
ATPase activity of the wildtype and mutant aqMutLs. (A) The ATPase activity was measured at 70°C by monitoring the release of phosphate ions with BIOMOL Green. The apparent rate constants were plotted against ATP concentrations. Average values from three independent experiments are shown with standard deviations. Theoretical Michaelis-Menten curves are overlaid. (B) The log*k*_cat_ values of the K28E, K287A, and K79R aqMutLs were plotted against pH. Dotted lines for the K28E and K287A aqMutLs are theoretical curves derived with p*K*_a_ values 9.2 and 9.3, respectively.

### Dimerization of the ATPase domain seems to be necessary for the ATPase activity of aqMutL

Around the ATP-binding site in the crystal structure of the aqMutL NTD, there is no other amino acid residue whose side chain is expected to have an alkaline p*K*_a_ value (Fig. 3A). Then, we suspected that, in the catalytic process of the ATPase reaction, unstructured loop regions of aqMutL might transform into the ordered structures just like the ATP-bound state of *E. coli* MutL. In *E. coli* MutL, the ATP binding induces transformation of the unstructured loop regions into the dimeric interface, in which Arg95 and Asn302 are included (23,43). The R95F/N302A double mutation has been reported to decrease the ATPase activity of *E. coli* MutL by 60 fold: an evidence that the transformation of the loop regions and the resulting dimerization of the ATPase domain are essential for the ATP hydrolysis by *E. coli* MutL. Our previous study revealed that the R95W mutation had no effect on the ATPase activity of aqMutL (36). Then, we examined the effect of the substitution of an alanine for Asn282 of aqMutL that corresponds to Asn302 of *E. coli* MutL. The CD measurements of the N282A aqMutL showed that the mutation does not influence the structure and stability of monomer aqMutL (Fig. 4A and B). The *k*_cat_ value of the N282A aqMutL was 52-fold smaller than that of the wildtype aqMutL (Fig. 5A and Table 1), indicating that the dimeric structure of the ATPase domain is also transiently formed in aqMutL during the catalysis.

We constructed a model structure of the ATP-bound dimeric form of the aqMutL NTD by using the ATP analog-bound state of the *E. coli* MutL NTD as a template structure for homology modeling. A single subunit of the resulting dimer is shown in Fig 3B. In the model structure, the side chains of Lys79 and Lys287 interacted with the β- and γ-phosphoryl groups of ATP, respectively. The interaction of Lys79 with the β-phosphoryl group of ATP has already been mentioned in a previous study (23). Both Lys79 and Lys287 are not included in the Bergerat ATP-binding fold, but conserved in all MutL proteins (Fig. 1). As for Lys287, the corresponding residue Lys307 of *E. coli* MutL has been shown to be involved in the ATPase activity; the *k*_cat_ value of the K307A *E. coli* MutL is 20-fold smaller than that of the wildtype (23). The lysine residue corresponding to Lys79 of aqMutL has not been investigated in *in vitro* experiments with MutLs from any species. Previously, a genetic study reported that the substitution of a glutamate for this lysine residue resulted in the loss of *in vivo* MMR activity in *Bacillus subtilis* (44). It was also shown that the mutation resulted in the absence of MutL protein in the *B. subtilis* cell. Therefore, involvement of the lysine residue in the ATPase hydrolysis has been unclear. It should be noted that Lynch syndrome-associated mutations K91R and K84E, the corresponding lysine residues of human PMS2 and MLH1, respectively, have been clinically described and classified as VUSs.

### Not Lys287 but Lys79 is the general acid catalyst for the ATPase activity of aqMutL

We prepared the K79A and K287A aqMutLs. Both mutant aqMutLs were properly folded and as stable against heat as the wildtype protein (Fig. 4A and B). The K287A aqMutL retained the ATPase activity; the mutation reduced the *k*_cat_ value by 3.3 fold with little effect on the *K*_m_ value (Fig. 5A and Table 1). The pH profile of the *k*_cat_ values of the K287A aqMutL exhibited the p*K*_a_ value of 9.3 (Fig. 5B), indicating that Lys287 is not the general acid. Since the ε-amino group of Lys287 is expected to interact with the γ-phosphoryl group of ATP (Fig. 3B), Lys287 might act as a Lewis acid that provides the positive charge to stabilize a negatively charged transient intermediate. On the other hand, the K79A mutation resulted in a complete loss of the ATPase activity (Fig. 5A). The loss of the activity was also observed when the general base Glu29 was replaced by an alanine (Fig. 5A).

In order to exclude the possibility that Lys79 is involved in the ATP binding, not the catalysis, trypsin limited proteolysis of the K79A aqMutL NTD was performed in the presence of an ATP analog, AMPPNP. It was previously shown that the ATP binding induced conformational changes of the aqMutL NTD, resulting in alteration of the digestion pattern by trypsin (36). As shown in Fig. S2A, the digestion pattern of the E29A aqMutL NTD, of which the ATP-binding ability was significantly reduced (23,36), was not influenced by the presence of AMPPNP. On the other hand, the digestion pattern of the K79A aqMutL NTD was almost the same as that of the wildtype NTD. The ATP-binding ability of the K79A aqMutL NTD was also examined by equilibrium dialysis (Fig. S2B), which revealed that the *K*_d_ values of the wildtype and K79A aqMutL NTDs for AMPPNP were 0.30 mM and 0.34 mM, respectively. These results indicate that Lys79 is not involved in the ATP binding.

**Supplementary Figure S2.**
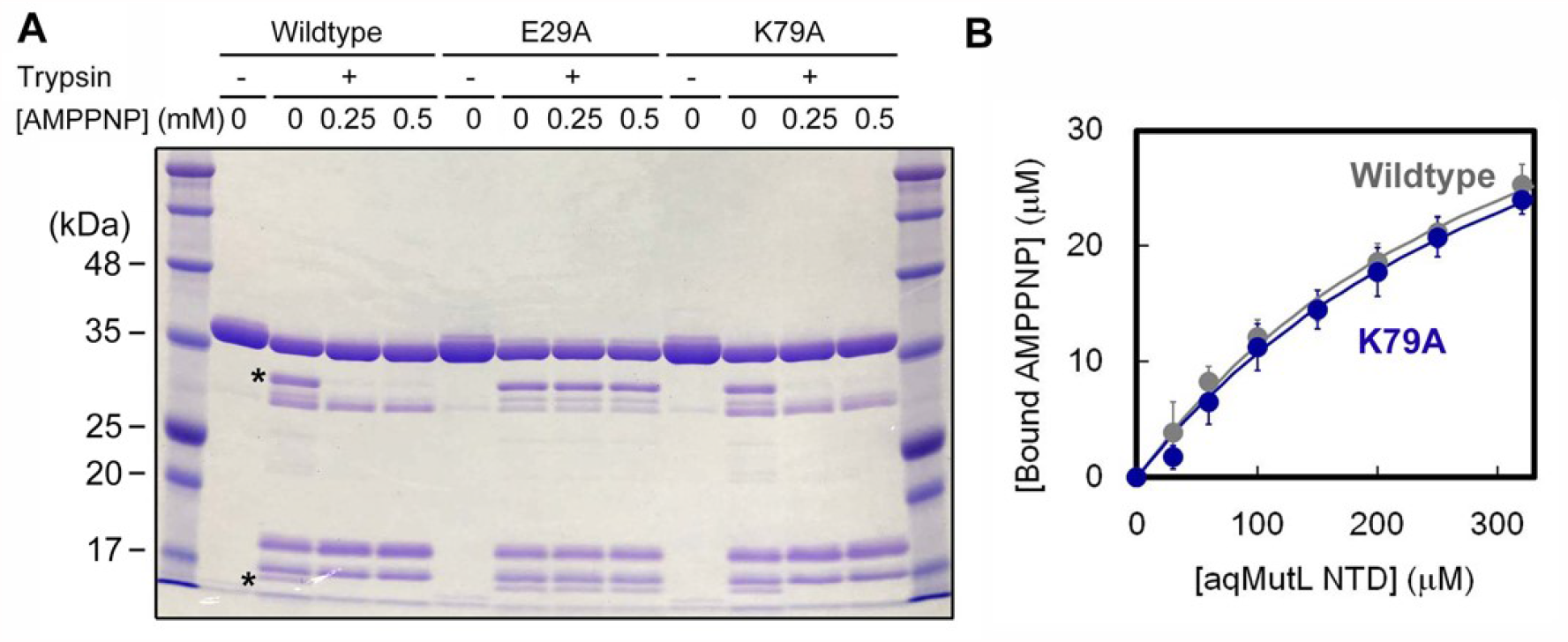
The K79A aqMutL NTD retained the ATP-binding ability. (A) Trypsin limited proteolysis of the wildtype, E29A, and K79A aqMutL NTDs was performed in the presence of 0, 0.25, and 0.5 mM AMPPNP. In the experiments with the wildtype and K79A aqMutL NTDs, the presence of AMPPNP prevented the generation of fragments indicated by asterisks. (B) Binding affinity was quantified by equilibrium dialysis. For each protein concentration, the concentration of unbound AMPPNP was determined on the basis of the absorbance at 260 nm of the solution from the buffer chamber after equilibrium was attained (see Materials and Methods). The concentrations of bound AMPPNP were calculated by subtracting those of unbound AMPPNP from the total concentration, and plotted against the protein concentrations. Average values from three independent experiments are shown with standard deviations. Theoretical curves with the *K*_d_ values of 0.30 mM and 0.34 mM for the wildtype and K79A aqMutL NTDs, respectively, are overlaid.

To test whether Lys79 is the general acid, we created the K79R aqMutL and examined its physicochemical properties and ATPase activity. The CD measurements revealed that the K79R mutation did not influence the structure and stability of the protein (Fig. 4A and B). As shown in Fig. 5A and Table 1, K79R retained the ATPase activity, although the mutation resulted in an 8.8-fold decrease in the *k*_cat_ value. Importantly, the *k*_cat_ values of the K79R aqMutL exhibited no pH dependency in the alkaline pH region (Fig. 5B). The p*K*_a_ of the guanidino group of arginine (12.5 in its free amino acid form) is much higher than that of the ε-amino group of lysine (10.0). The apparent disappearance of the p*K*_a_ in the pH profile would thus be a consequence of the alkaline shift of the pH dependence of the activity. These findings indicate that Lys79 is the general acid catalyst and that the arginine residue at the position 79 is not optimal but still functional as the general acid.

### The lysine residues corresponding to aqMutL Lys79 are also essential for the ATPase activity of human PMS2 and MLH1

Lys79 and Lys287 of aqMutL correspond to Lys91 and Lys340, respectively, in human PMS2. In order to verify the importance of these residues in the ATPase activity, the wildtype and mutant forms of the human PMS2 NTD were prepared. The K340A and K340E PMS2 NTDs exhibited weakened, but substantial, ATPase activities, whose *k*_cat_ values were 5.9- and 13-fold smaller than that of the wildtype (Fig. 6A and Table 1). In contrast, the K91A mutation completely abolished the ATPase activity of the human PMS2 NTD (Fig. 6A). In addition, the K91R PMS2 NTD retained the ATPase activity, although the *k*_cat_ value was decreased by 8.4 fold (Fig. 6A and Table 1).

**Figure 6.**
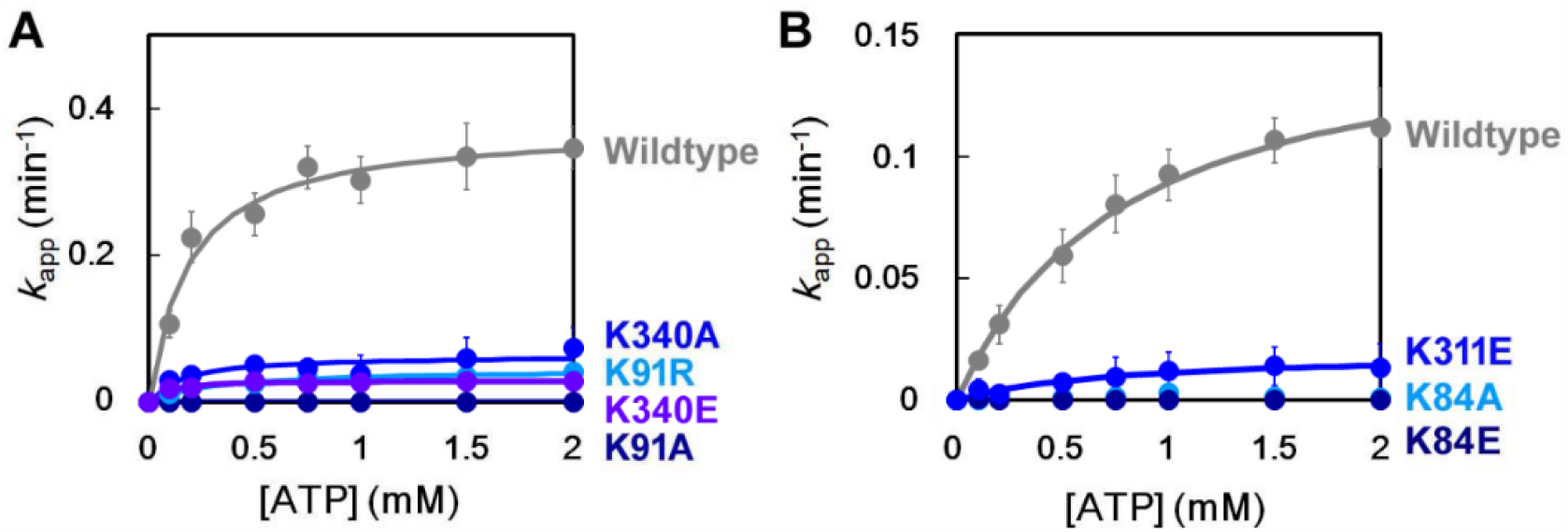
ATPase activity of the wildtype and mutant ProS2-fused human PMS2 NTDs (A) and histidine-tagged human MLH1 NTDs (B). The K91R and K340E mutations are clinically found in human PMS2. Similarly, K84E and K311E are the clinically-found mutations of human MLH1. The ATPase activity was measured at 37°C. The average values from three independent experiments are shown with standard deviations. Theoretical Michaelis-Menten curves are overlaid.

The same results were obtained for the human MLH1 NTD. Lys79 and Lys287 of aqMutL correspond to Lys84 and Lys311, respectively, in human MLH1. As shown in Fig. 6B and Table 1, the K311E MLH1 NTD retained a weak ATPase activity; the *k*_cat_ value was 8.0-fold smaller than that of the wildtype. On the other hand, the activity was completely abolished by the K84A and K84E mutations; the latter mutation was found in Lynch syndrome-suspected patients and diminished the MMR activity in the human cell lysate without decrease in the protein expression (45). Since human MutL homologs were not stable at alkaline pHs, the pH profile of the ATPase activity could not be examined.

The results for these mutant forms of aqMutL, the human PMS2 NTD, and the human MLH1 NTD were consistent with each other, which strongly supports the notion that the lysine residue corresponding to Lys79 of aqMutL serves as the general acid in the MutL reaction.

As for another lysine residue, the recombinant K40E PMS2 NTD, which corresponds to the K28E aqMutL, was degraded in the host *E. coli* cell probably due to the mutation-induced destabilization of the protein structure. Similarly, the K33E MLH1 NTD precipitated during overexpression in the host cell. These are in agreement with the result that K28E mutation in aqMutL resulted in the decreased protein stability (Fig. 4B and 4C). Previously, we reported that another Lynch syndrome-associated mutation S34I of aqMutL (S46I of human PMS2), which is at the Bergerat ATP-binding fold, also reduces stability of the overall protein structure through destabilization of a local structure around the ATP-binding site (36). Integrity of the Bergerat ATP-binding fold would be essential for the structural stability of MutL.

### The ATPase activity-related lysine residues are essential for MMR in vivo

As described above, three lysine residues, Lys28, Lys79, and Lys287, are involved in the *in vitro* ATPase activity. Importance of the three lysine residues in the MMR activity in the cell was examined by the phenotype complementation experiments using a eubacterium, *Thermus thermophilus*, one of the model organisms that has been utilized for studies on DNA repair including MMR (46,47). Dysfunction of MMR causes a significant increase in spontaneous mutation frequency that can be measured as the frequency of streptomycin-resistant bacterial cells (47). As shown in Fig. 7, the frequency of streptomycin-resistant cells of the *mutL*-lacking *T. thermophilus* strain was approximately 10-fold higher than that of the wildtype strain.

**Figure 7.**
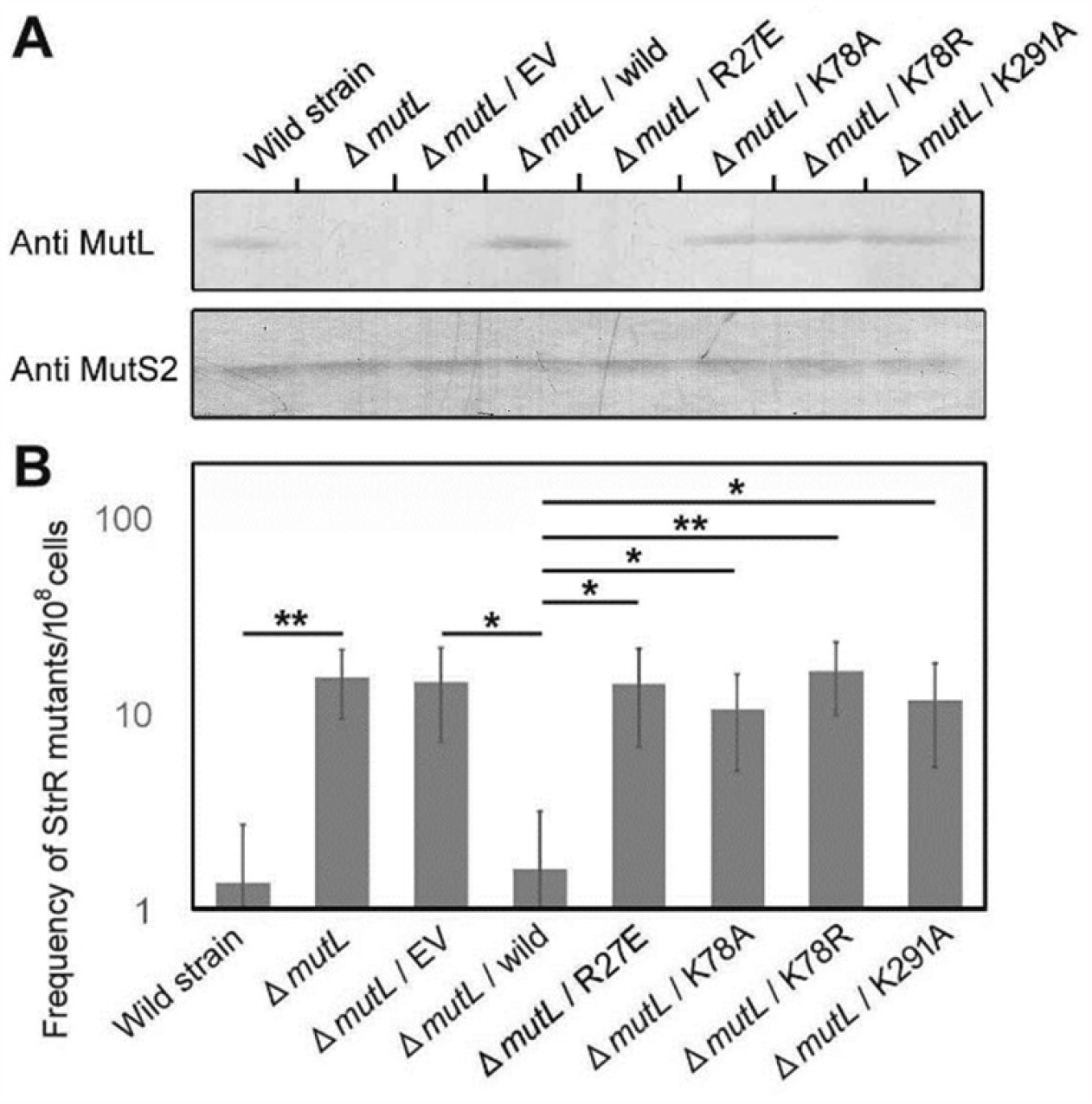
Phenotype complementation assay using *T. thermophilus*. (A) Protein expression in *T. thermophilus* cells was detected by Western blot analysis using anti-*T. thermophilus* MutL and anti-*T. thermophilus* MutS2 antisera. (B) Frequency of spontaneous mutagenesis was evaluated by measuring the frequency of streptomycin-resistant mutants. EV means the empty vector. The results are shown as averages of five independent experiments with standard deviations. Statistical analyses were performed with the two-sided Student’s *t* test: *, *p* < 0.05; **, *p* < 0.01.

Western blot analysis showed that introduction of the wildtype *T. thermophilus mutL* gene led to successful expression of the gene in the *mutL*-lacking strain (Fig. 7A). Expression of the wildtype *mutL* gene reduced the mutability of the *mutL*-lacking strain to a level equal to that of the wildtype strain (Fig. 7B). In contrast, the *mutL* gene carrying the R27E mutation, which corresponds to the K28E mutation of aqMutL, did not complement the hyper mutator phenotype of the disruptant (Fig. 7B). Interestingly, Western blot analysis revealed that the R27E *T. thermophilus* MutL was absent (Fig. 7A) probably due to degradation in the cell. This is consistent with the result that the K28E aqMutL was unstable compared with the wildtype protein. Genes for the K78A, K78R, and K291A *T. thermophilus* MutLs, which correspond to the K79A, K79R, and K287A aqMutLs, were also introduced into the *mutL*-lacking *T. thermophilus*. Although Western blot analysis confirmed the successful expression of these variant genes in the cell (Fig 7A), these did not complement the hyper mutator phenotype of the *mutL*-lacking strain (Fig. 7B). These results indicate that the lysine residues are critical for *in vivo* MMR and that the reduction of the *in vitro* ATPase activity by only 3 fold results in almost loss of the *in vivo* MMR activity.

### Catalytic mechanism of the ATPase activity of MutL

Based on the results presented above, we propose a catalytic mechanism of the MutL ATPase activity (Fig. 8). Since phosphorus is able to form trivalent, tetravalent, and pentavalent compounds, a phosphoryl transfer reaction can proceed by either of associative and dissociative mechanisms. In the dissociative mechanism, the γ-phosphate of ATP is first eliminated to generate a metaphosphate ion (PO_3_ ^-^) and ADP, and, then, a nucleophile (HO^-^) activated by a general base is added to the metaphosphate. In previous reports, *T. thermophilus* and *E. coli* MutLs carrying mutations at the general base Glu29 were not able to generate the product ADP (23,48). These support that the ATP hydrolysis by MutL does not proceed by the dissociative mechanism. Therefore, we propose the following catalytic mechanism based on the associative mechanism: The general base Glu29 of aqMutL orients and activates the nucleophilic water molecule. The nucleophilic attack of the water molecule to the γ-phosphoryl group of ATP generates a transient pentacovalent intermediate, which is stabilized by the positive charge of the ε-amino group of the Lewis acid Lys287. Subsequently, Lys79, the general acid catalyst, donates a proton to the β-phosphoryl group to complete the cleavage reaction.

**Figure 8.**
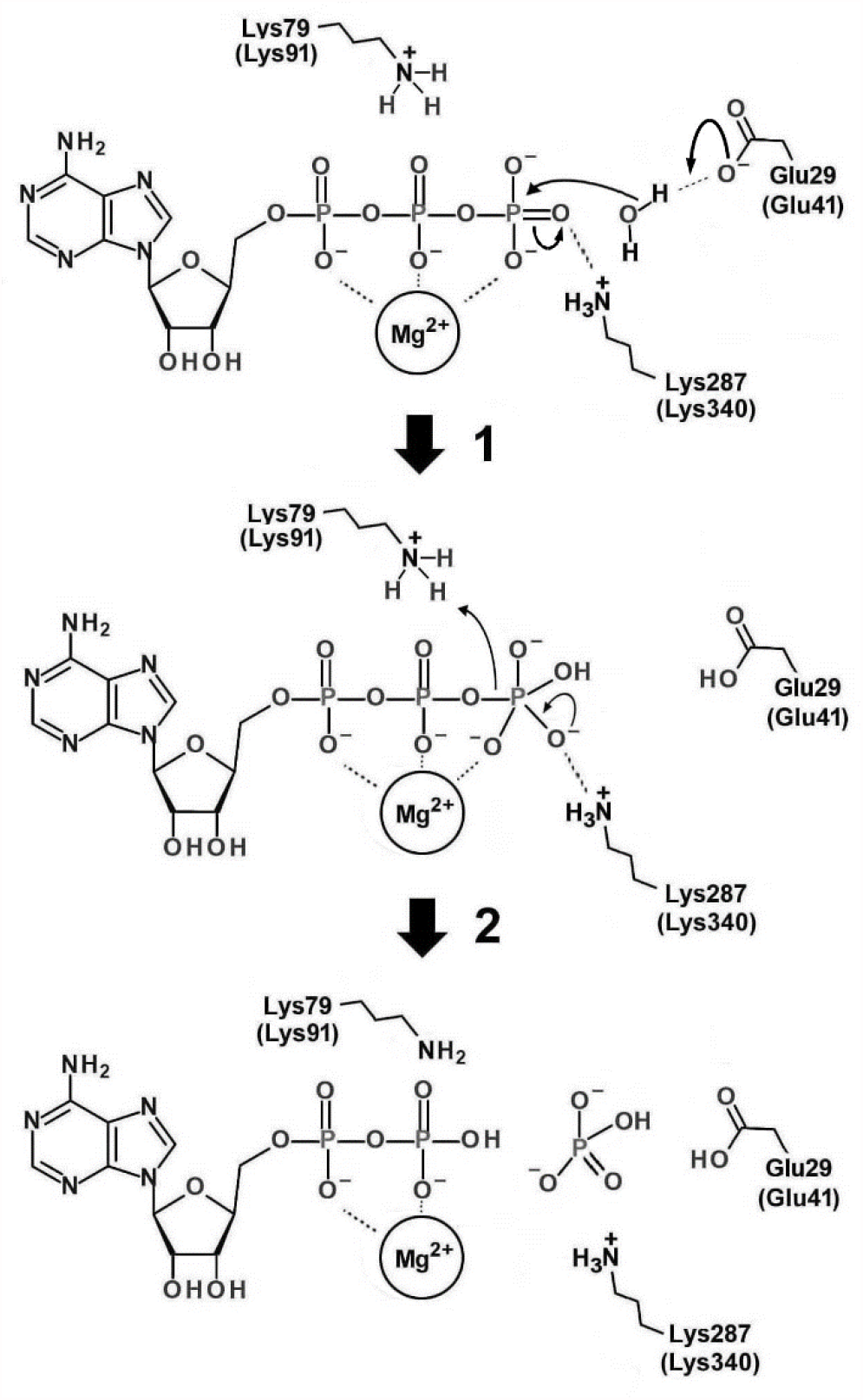
A model for the catalytic mechanism of the MutL ATPase reaction. Numbering of the amino acid residues is based on the numbering of the aqMutL sequence. Numbering of the human PMS2 is indicated in the parentheses. ATP and magnesium ion are bound at the ATPase active site of MutL. Step 1: The nucleophilic water molecule is coordinated and activated by the general base Glu29. Attack of the nucleophilic water to the γ-phosphoryl group generates a transient pentacovalent intermediate with an extra negative charge, which is stabilized by the positive charge from Lys287. Step 2: Instability of the negative charge on the phosphoryl oxygen leads to re-formation of a double bond and collapse of the intermediate, breaking the phosphoanhydride bond between the β- and γ-phosphoryl groups. The general acid Lys79 provides a proton to the β-phosphoryl group, facilitating the bond breakage.

Glu29, Lys79, and Lys287 of aqMutL correspond to Glu42, Lys103, and Lys337, respectively, of *E. coli* GyrB, another GHKL superfamily ATPase (Fig. S3A–C). It has been considered that, in the ATPase reaction of *E. coli* GyrB, Glu42 serves as a general base to activate the nucleophilic water molecule and that Lys337 stabilizes the negatively-charged intermediate (49,50). It was previously reported that substitutions of Ile, Thr, or Glu for Lys103 of *E. coli* GyrB completely abolished the ATPase activity (51), although another study reported that the K103I and K103A *E. coli* GyrBs retained slightly-weakened ATPase activities (52). We assessed the effect of the Ala substitution at the corresponding Lys residue (Lys109) on the ATPase activity of the *A. aeolicus* GyrB (aqGyrB) NTD. As shown in Fig. S3D and Table 1, the ATPase activity of the L109A aqGyrB NTD was below the detection limit, indicating that the Lys residue also acts as a general acid in aqGyrB.

**Supplementary Figure S3.**
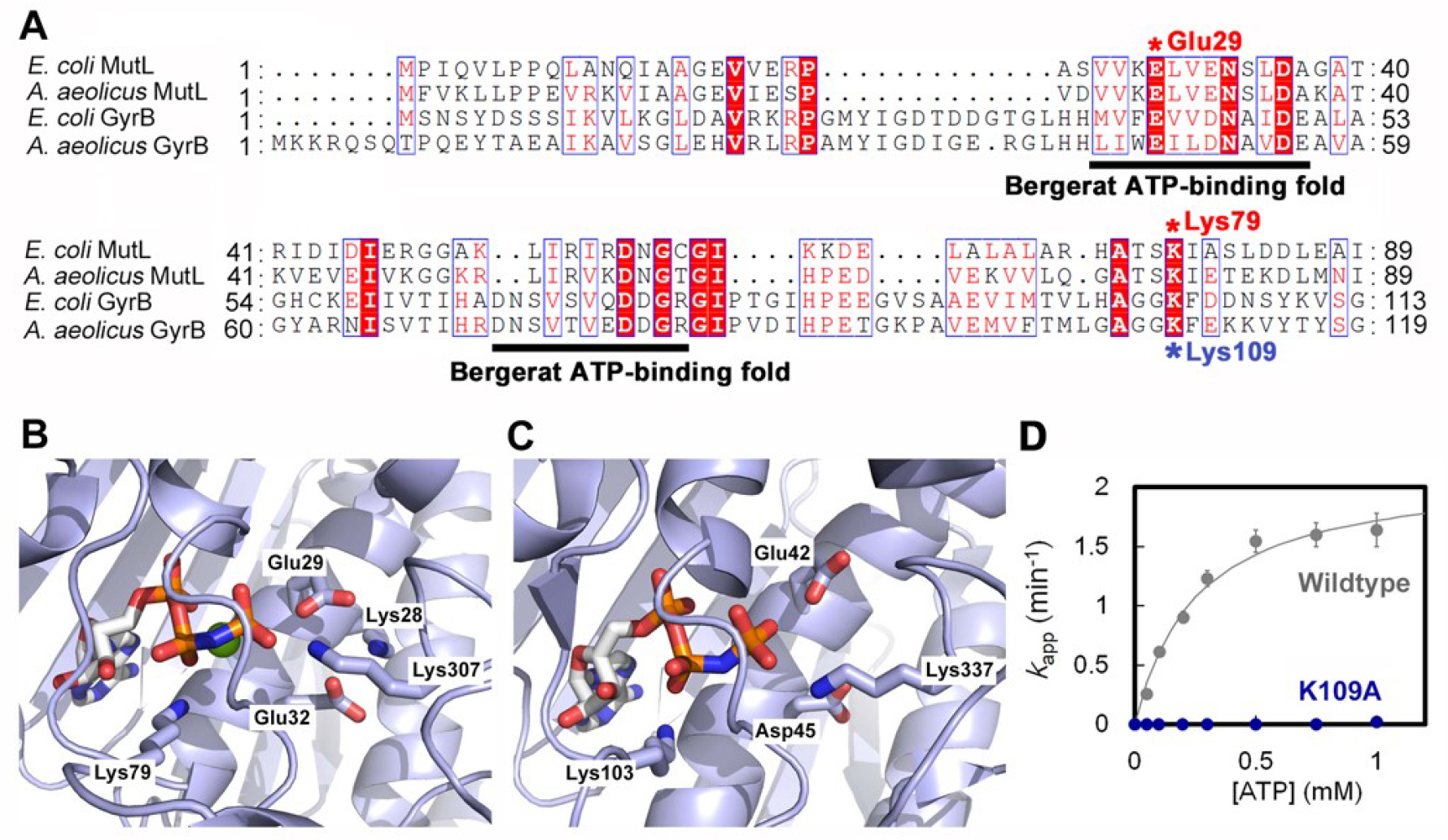
ATPase activity of GyrB. (A) Amino acid sequence alignment of MutL and GyrB. The amino acid sequences of the N-terminal regions of *E. coli* MutL, aqMutL, *E. coli* GyrB, and aqGyrB were aligned by CLUSTALW and visualized by using the ESPript3 program. Red asterisks indicate the general base catalyst (Glu29) and the putative general acid catalyst (Lys79) of *E. coli* MutL. A blue asterisk indicates the putative general acid catalyst (Lys109) of aqGyrB. The ATPase catalytic site of *E. coli* MutL (PDB ID: 1B63) (B) and that of *E. coli* GyrB (PDB ID: 1EI1) (C). (D) The ATPase assay of the wildtype and K109A aqGyrB NTDs. Average values from three independent experiments are shown with standard deviations. The theoretical Michaelis-Menten curve for the wildtype aqGyrB NTD is overlaid.

### Pathogenicity of the Lynch syndrome-associated VUSs

Various Lynch syndrome-associated mutations of human PMS2 and MLH1 have been found at the lysine residues, of which the involvement in the ATPase activity was revealed by the present study. The pathogenicity of the K91R and K340E mutations of human PMS2 can be explained by reduction in the ATPase activity, and so be the mutations K84E and K311E of human MLH1. The K40E and K33E mutations of human PMS2 and MLH1, respectively, would be pathogenic because these mutations destabilize the protein structure. Thus, pathogenicity of these previously-found VUSs in MutL homologs can now be clearly explained.

## Materials and Methods

### Construction of expression plasmids

The expression plasmids for the K28E, E29A, K79A, K79R, N282A, and K287A full-length aqMutLs were constructed by introducing the mutations into the pET-11a/aq*mutL* plasmid (47) using the PrimeSTAR mutagenesis procedure (Takara). The primer sets used for the construction of these mutant aqMutLs were 5’-TCGTAGAGGAACTCGTTGAGAATTCT-3’ and 5’-CGAGTTCCTCTACGACGTCAACAGGA-3’, 5’-GTAAAGGCGCTCGTTGAGAATTCTCTG-3’ and 5’-AACGAGCGCCTTTACGACGTCAACAGG-3’, 5’-ACGAGTGCGATAGAGACCGAAAAGGA-3’ and 5’-CTCTATCGCACTCGTCGCTCCCTGCAG-3’, 5’-ACGAGTAGGATAGAGACCGAAAAGGA-3’ and 5’ CTCTATCCTACTCGTCGCTCCCTGCA-3’, 5’-GACTTCGCCGTCCACCCCAAGAAAAA-3’ and 5’-GTGGACGGCGAAGTCAACCATAAAGGG-3’, and 5’-CCCAAGGCAAAGGAAGTGAACATACT-3’ and 5’-TTCCTTTGCCTTGGGGTGGACGTTGAA-3’, respectively.

The expression plasmids for the K40E, K91A, K91R, K340A, and K340E human PMS2 NTDs (residues 1–365) were constructed by introducing the mutations into the pCold ProS2/human *PMS2 NTD* plasmid (36) using the PrimeSTAR mutagenesis procedure. The primer sets used were 5’-GCGGTAGAGGAGTTAGTAGAAAACAG-3’ and 5’-TAACTCCTCTACCGCAGTGCTTAGACT-3’, 5’-CATCTGCGATTCAAGAGTTTGCCGA-3’ and 5’-TTGAATCGCAGATGTGTGATGTTTCA-3’, 5’-CATCTAGGATTCAAGAGTTTGCCGA-3’ and 5’-TTGAATCCTAGATGTGTGATGTTTCA-3’, 5’-TCCAGATGCAAGGCAAATTTTGCTACA-3’ and 5’-TGCCTTGCATCTGGAGTAACATTGAT-3’, and 5’-CCAGATGAAAGGCAAATTTTGCTAC-3’ and 5’-TGCCTTTCATCTGGAGTAACATTGAT-3’, respectively.

The cDNA of human *MLH1* was obtained from the National Institute of Technology and Evaluation (Kisarazu, Japan). The nucleotide sequence corresponding to the NTD of human MLH1 (residues 1–337) was amplified by PCR using the primer set 5’-ATAT<u>CATATG</u>TCGTTCGTGGCAGGG-3’ and 5’-TAT<u>CTCGAG</u>TTAGCCCAGGAGCTTGCTCTC-3’ (underlines indicate NdeI and XhoI sites, respectively). The amplified fragment was digested with NdeI and XhoI and ligated into the corresponding sites of the pET28-MHL (37) to obtain the pET28-MHL/human *MLH1 NTD*. The expression plasmids for the K33E, K84E, K84A, and K311E human MLH1 NTD were constructed by introducing the mutations into the pET28-MHL/human *MLH1 NTD* plasmid using the PrimeSTAR mutagenesis procedure. The primer sets used were 5’-TGCTATCGAAGAGATGATTGAGAACT-3’ and 5’-ATCTCTTCGATAGCATTAGCTGGCCG-3’, 5’-TACTAGTGAACTGCAGTCCTTTGAGG-3’ and 5’-TGCAGTTCACTAGTAGTGAACCTTTC-3’, 5’-TACTAGTGCACTGCAGTCCTTTGAGG-3’ and 5’-TGCAGTGCACTAGTAGTGAACCTTTC-3’, and 5’-CCCCACAGAGCATGAAGTTCACTTCC-3’ and 5’-TCATGCTCTGTGGGGTGCACATTAAC-3’.

The nucleotide sequence encoding the NTD (residues 1–397) of aqGyrB was amplified by PCR using the *A. aeolicus* genomic DNA as a template. The following primers were used: 5’-ATA<u>CATATG</u>AAGAAAAGGCAGTCTCAAACC-3’ and 5’-AT<u>GGATCC</u>TTAAACGAGTTCCTTAGCTTTTTTTGC-3’ (underlines indicate NdeI and BamHI sites, respectively). The amplified DNA fragment was digested with NdeI and BamHI and ligated into the corresponding sites of the pET-11a (Novagen) to obtain the pET-11a/*aqgyrB NTD*. The expression plasmid for the K109A mutant form of the aqGyrB NTD was constructed by introducing the mutation into the pET-11a/*aqgyrB NTD* plasmid using the PrimeSTAR mutagenesis procedure with the primers: 5’-GGAGGAGCGTTTGAAAAGAAAGTCTA-3’ and 5’-TTCAAACGCTCCTCCCGCCCCGAGCAT-3’.

Sequence analyses revealed that, apart from the desired mutations, no unintended mutation was introduced into the *A. aeolicus mutL, gyrB*, human *PMS2* and *MLH1* genes.

### Expression and purification of proteins

The histidine-tagged human MLH1 NTD was prepared using the modified procedure of the previously-described method (37). Rosetta2(DE3) pLysS *E. coli* competent cells were transformed with the pET28-MHL/human *MLH1 NTD* carrying the wildtype, K33E, K84A, K84E, or K311E NTD gene. One liter of LB medium (Difco) containing 50 μg/ml kanamycin and 30 μg/ml chloramphenicol was inoculated with 5 ml of overnight preculture. After 4 h of cultivation at 37°C, isopropyl-β-D-thiogalactopyranoside was added to a final concentration of 0.2 mM. The cells were further cultivated at 15°C for 24 h and harvested by centrifugation.

The cells were re-suspended with 20 ml of 50 mM Tris-HCl (pH 8.0) containing 500 mM NaCl, 5% glycerol, and 0.5% CHAPS, and disrupted by sonication on ice. After centrifugation at 48,000 × g for 20 min, the supernatant was loaded onto a His-bind resin column (Merck Millipore) (1 ml) pre-equilibrated with buffer I containing 20 mM Tris-HCl (pH 8.0), 5% glycerol, and 500 mM NaCl. The column was washed with 2 ml of buffer I and further washed with 10 ml of buffer I containing 50 mM imidazole. The column was eluted with 6 ml of buffer I containing 100 mM EDTA. The fractions containing the histidine tagged human MLH1 NTD were collected and dialyzed against 50 mM Tris-HCl (pH 8.0), 5 % glycerol, and 250 mM NaCl. The protein solution was loaded onto a Toyopearl SP-650 column (TOSOH) (10 ml) pre-equilibrated with 20 mM Tris-HCl (pH 8.0) containing 20 mM NaCl. The column was washed with 50 ml of 20 mM Tris-HCl (pH 8.0) containing 100 mM NaCl. The column was eluted with 15 ml of 20 mM Tris-HCl (pH 8.0) containing 250 mM NaCl. The protein solution was concentrated and stored at 4°C.

The wildtype and mutant forms of the full-length aqMutL and the aqMutL NTD were overexpressed in *E. coli* and purified as previously described (36,47). The wildtype and mutant forms of the human ProS2-fused PMS2 NTD were also overexpressed in *E. coli* and purified by the previously-described procedure (36).

The wildtype and K109A aqGyrB NTDs were overexpressed in *E. coli* by the same procedure as that for aqMutL. The cells were lysed and heat-treated at 70°C for 10 min and immediately chilled on ice for 10 min. After centrifugation at 48,000 × g for 20 min, the supernatant was loaded onto a Toyopearl SP column (20 ml) (TOSOH) pre-equilibrated with 30 mM Tris-HCl (pH 8.0). The column was washed with 30 ml of 30 mM Tris-HCl (pH 8.0) and further washed with 60 ml of 30 mM Tris-HCl (pH 8.0) containing 0.5 M NaCl. The column was eluted with 60 ml of 30 mM Tris-HCl (pH 8.0) containing 1.0M NaCl. The fractions containing aqGyrB NTD were detected by SDS-PAGE and collected. Ammonium sulfate was added to the fractions to yield a final concentration of 1 M. The solution was loaded onto a Toyopearl Phenyl column (10 ml) (TOSOH) equilibrated with 30 mM Tris-HCl (pH 8.0) containing 1 M ammonium sulfate. The column was washed with 30 ml of 30 mM Tris-HCl (pH 8.0) containing 1 M ammonium sulfate and further washed with 30 ml of 30 mM Tris-HCl (pH 8.0) containing 0.3 M ammonium sulfate. Then, the column was eluted with 50 ml of 30 mM Tris-HCl (pH 8.0). The fractions containing the protein (30 ml) were collected and dialyzed against one liter of 30 mM Tris-HCl (pH 8.0) three times at 4°C. The protein solution was concentrated and stored at 4°C.

### CD spectrometry

CD measurements were performed with the Jasco spectropolarimeter, model J-720W (Jasco). Measurements of CD spectra were carried out in a solution comprised of 50 mM potassium phosphate (pH 7.0) and ∼10 μM protein using a 0.1-cm cell at 25°C. The residue molar ellipticity [θ] was defined as 100θ_obs_/(*lc*), where θ_obs_ is the observed ellipticity, *l* is the length of the light path in centimeters, and *c* is the residue molar concentration of the protein. The temperature scans were carried out with a heating rate of 1°C/min controlled by the Jasco programmable Peltier element.

Urea-dependent denaturation of aqMutL was examined by measuring the intensity of CD at 222 nm after incubation of aqMutL with various concentrations of urea at room temperature for 16 h.

For the measurements of CD spectra under different pH conditions, 30 mM sodium acetate, 30 mM MES-NaOH, 30 mM Tris-HCl, or 30 mM sodium carbonate/bicarbonate was used to cover the pH range from 3.5 to 5.5, from 6.5 to 7.0, from 7.5 to 9.0, or from 9.2 to 10.7, respectively. After incubation of ∼10 μM protein at 60°C for 3 h, CD spectra from 200 to 250 nm were measured.

### ATPase assay

The ATPase activity of aqMutL, the histidine-tagged human MLH1 NTD, and the ProS2-fused human PMS2 NTD was assayed by the previously described procedure (36,53). The assay was carried out with 10 μM protein in a 20-μL reaction mixture comprised of 50 mM HEPES-KOH (pH 7.5), 100 mM NaCl, 5 mM MgCl_2_ and various concentrations of ATP. The reactions were performed at 70°C for 60 min with aqMutL and the aqGyrB NTD. In the assay for the NTDs of human MutL homologs, reactions were performed at 37°C for 120 min. The reactions were stopped by the addition of 200 μL of BIOMOL Green reagent (Enzo Life Sciences). Mixtures were incubated at room temperature for 30 min to allow the color development. The absorbance at 620 nm was measured by a microplate reader 680XR (Bio-Rad Laboratories). The experiments were performed in triplicate. The standard Michaelis-Menten equation was fitted to the data to determine the *k*_cat_ and *K*_m_ values.

For the measurements of the ATPase activity of aqMutL under different pH conditions, reactions were performed at 60°C, and in place of 50 mM HEPES-KOH, 50 mM sodium acetate, 50 mM MES-NaOH, 50 mM Tris-HCl, or 50 mM sodium carbonate/bicarbonate was used to cover the pH range from 3.5 to 5.5, from 6.5 to 7.0, from 7.5 to 9.0, or from 9.2 to 10.7, respectively. Measurements at pH 5.5, 7.5, and 9.0 were carried out with two kinds of buffering agents, sodium acetate or MES-KOH for pH 5.5, MES-KOH or Tris-HCl for pH 7.5, and Tris-HCl or sodium carbonate/bicarbonate for pH 9.0, which revealed that difference of the agents did not affect the ATPase activity of aqMutL.

The pH profile of the aqMutL ATPase activity was analyzed as follows: In the presence of saturating amounts of the substrate, all of the enzyme molecules in the reaction mixture are essentially in the form of the enzyme-substrate complex (ES). Accordingly, the pH profile can be analyzed by the simple scheme:

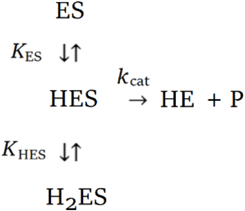

where ES, HES, and H_2_ES are unprotonated, singly-protonated, and doubly-protonated ES, respectively, and only HES is enzymatically active; *K*_ES_ and *K*_HES_ are the acid dissociation constants for the corresponding protonation/deprotonation steps. The *k*_cat_ value at each pH ((*k*_cat_)_H_) is defined by

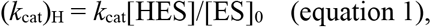

where [ES]_0_ is the total concentration of the enzyme-substrate complex and defined by [ES]_0_ = [ES] + [HES] + [H_2_ES]. The dissociation constants are defined by *K*_HES_ = [H^+^][HES]/[H_2_ES] and *K*_ES_ = [H^+^][ES]/[HES].

From these,

[ES]_0_ = [HES] ([H^+^]/*K*_HES_ + 1 + *K*_ES_/[H^+^]), and

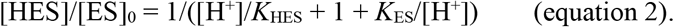

By substituting the equation 2 for [HES]/[ES]_0_ in the equation 1, the pH dependence of *k*_cat_ is obtained:

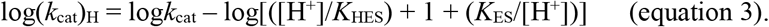

p*K*_HES_ and p*K*_ES_ values were obtained by fitting the equation 3 to the data using Igor Pro 8.

### Complementation experiment

The *mutL*-lacking *T. thermophilus* HB8 strain was constructed as previously described (47). The wildtype, R27E, K78A, K78R, and K291A *T. thermophilus mutL* genes were ligated into the pMK18::HygΔKan plasmid (54,55) using XbaI and HindIII restriction sites. These plasmids were then transformed into the *mutL*-lacking *T. thermophilus* HB8 strain. The frequency of streptomycin-resistant cells of the transformants were measured by the previously described method (47).

### Western Blotting

*T. thermophilus* HB8 cells at the late-exponential growth phase were harvested by centrifugation and suspended in a buffer containing 50 mM Tris-HCl (pH 8.0), 100 mM NaCl, 10% glycerol.

The cells were lysed by ultrasonication on ice. After centrifugation, total protein concentrations in the supernatants were determined by Bradford method. Ten μg total protein of each sample was subjected to SDS-PAGE (10 % polyacrylamide gel) and electroblotted onto a nitrocellurose membrane. The membrane was incubated in a blocking solution comprising 20 mM Tris-HCl (pH 7.5), 500 mM NaCl, and 3% bovine serum albumin (nacalai) for 30 min at room temperature. After washing with the washing solution comprising 20 mM Tris-HCl (pH 7.5), 500 mM NaCl, and 0.05% Tween-20 (nacalai), the membrane was immersed into the blocking solution containing rabbit anti-*T. thermophilus* MutL or anti-*T. thermophilus* MutS2 antiserum and incubated for 1 h at room temperature. The membrane was washed with the washing solution and then, reacted with Protein A-horseradish peroxidase conjugate (Bio-Rad Laboratories) in the blocking solution for 2 h at room temperature. The membrane was washed with the washing solution twice and then reacted with 4-chloro-1-naphthol (Bio-Rad Laboratories) in the HRP color development buffer (Bio-Rad Laboratories) for 10 min at room temperature. The staining was stopped by washing the membrane in deionized water.

### Limited proteolysis

The wildtype and mutant forms of the aqMutL NTD (5 μM) were reacted with 30 ng/μl trypsin (from bovine pancreas; Sigma Aldrich) in a buffer consisting of 50 mM Tris-HCl (pH 8.0), 100 mM NaCl, 5 mM MgCl_2_, and 0, 0.25 or 0.5 mM AMPPNP at 37°C for 20 min. Reactions were stopped by addition of an equal volume of the SDS-containing dye comprised of 125 mM Tris-HCl (pH 7.5), 10% 2-mercaptoethanol, 4% SDS, 10% sucrose, and 0.01% bromophenol blue. Twenty microliter aliquot of each mixture was resolved by SDS-12.5% PAGE and stained with Coomassie brilliant blue.

### Equilibrium dialysis

Seventy five microliter of 50 mM Tris-HCl (pH 8.0), 100 mM NaCl, and 5 mM MgCl_2_ containing 50 μM AMPPNP was loaded into the buffer chamber of the DispoEquilibrium Dialyzer (10 kDa molecular weight cut off membrane) (Harvard Bioscience, Inc.). Subsequently, an equivalent volume of the aqMutL NTD (0–320 μM) in 50 mM Tris-HCl (pH 8.0), 100 mM NaCl, and 5 mM MgCl_2_ was placed into the opposite sample chamber of the device. The device was gently agitated on a shaker at room temperature for 16 h. The absorbance at 260 nm of the solution from the buffer chamber was measured. The concentration of unbound AMPPNP was calculated by using a molar extinction coefficient, 15,400 M^-1^ cm^-1^. The following equation was fitted to the data to determine *K*_d_ values of the wildtype and K79A aqMutL NTDs.

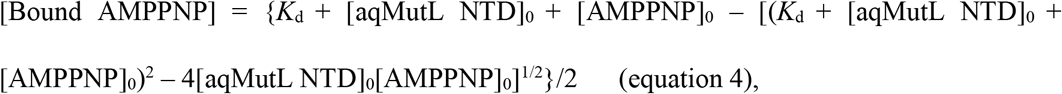

where [aqMutL NTD]_0_ and [AMPPNP]_0_ are the total concentrations of the protein and ligand, respectively.

## Data Availability

This study includes no data deposited in external repositories.

## Acknowledgements

This work was supported by JSPS KAKENHI grant number JP19K07376 (to KF). Authors thank Prof. Ryoji Masui (Osaka City University) for providing the pMK18::HygΔKan plasmid.

## Author contributions

K.F and T.Y conceptualized the project. K.F. performed all experiments. Y.F. constructed the18::HygΔKan plasmid. K.F. and T.Y. analyzed the data and wrote the manuscript with help from Y.F.

## Conflict of interest

The authors declare that they have no conflict of interest.

## Notes

### Competing Interest Statement

The authors have declared no competing interest.

